# A reduced glycosaminoglycan-linked residual-strain model captures regional opening angle changes after depletion in the porcine thoracic aorta

**DOI:** 10.64898/2026.07.15.738269

**Authors:** Michel Labrosse, Noor Ghadie, Jean-Philippe St-Pierre, Munir Boodhwani

## Abstract

Residual stresses in arteries are commonly revealed by the spring opening of a ring after a radial cut. Glycosaminoglycans (GAGs) contribute to this response, but fixed-charge-density (FCD)-driven Donnan swelling alone does not fully explain the opening-angle reduction measured after enzymatic GAG depletion. We therefore tested whether regional mean depletion responses are consistent with a removable preferred-stretch field shaped by the transmural FCD profile and superposed on a structural field retained after depletion.

A reduced analytical-computational axisymmetric closure framework was applied to regional-average measurements from the ascending aorta, arch, and descending porcine thoracic aorta. Each control state was fitted using its measured circumferential opening angle, geometry, material properties, and through-wall FCD profile. GAG depletion was represented by removing the FCD-linked preferred-stretch component. One coefficient governing this removable component was selected jointly from the three measured regional post-depletion angles. A one-layer wall was the primary parsimonious model; a two-layer wall tested anatomical robustness.

The one-layer model fitted a shared coefficient of −1.4226 *×* 10^−3^ (mEq*/*L)^−1^ and predicted depleted angles of 82.639°, 44.032°, and 19.979°, compared with measured values of 85°, 41°, and 18^°^ (three-region RMSE 2.50°). The two-layer model fitted −1.51746 *×* 10^−3^ (mEq*/*L)^−1^ and predicted 82.972°, 44.273°, and 18.534^°^ (RMSE 2.24°). Thus, layer differentiation improved aggregate fit only modestly.

These results show that the three regional mean depletion responses can be represented by one common FCD-linked removable preferred-stretch contribution. Because the model uses regional averages and fitted region-specific control structural fields, it does not establish specimen-level predictive validity or uniquely identify the underlying GAG-mediated mechanism. Donnan swelling remains mechanically relevant, but it is insufficient alone to explain the measured regional response.

**Statement of Significance:** Glycosaminoglycans are charged extracellular-matrix constituents that influence arterial residual stress through osmotic effects and interactions with the fibrous matrix. We tested whether the through-wall FCD pattern also defines a removable circumferential preferred-stretch component. One coefficient linking that pattern to the depletion response captured the regional mean post-depletion opening angles of the ascending aorta, arch, and descending aorta within 3.3° using a one-layer model. A two-layer model improved aggregate error only modestly. The result provides a compact mechanistic link between extracellular-matrix composition and residual opening without requiring a separately fitted GAG effect in each region.

## 1 Introduction

Residual stresses are a defining feature of arterial mechanics. When an unloaded arterial ring is cut radially, it springs open, revealing that the closed ring is not stress free [1–3]. Opening angle is therefore widely used as an indicator of circumferential residual stress and strain. These residual fields influence transmural stress under physiological loading, constitutive reference configurations, and interpretations of vascular growth and remodeling.

The aortic extracellular matrix contains elastin, collagen, and proteoglycans bearing GAG chains. GAGs carry fixed negative charge, generate Donnan osmotic swelling, and interact mechanically with the fibrous matrix [4]. Heterogeneous proteoglycan distributions can regulate aortic residual stress through swelling [5], while GAG removal also alters collagen-fiber waviness and recruitment [6, 7]. In porcine thoracic aorta, sulfated-GAG content varies through the wall and by region and is associated with opening angle [8]; enzymatic depletion directly reduces the opening angle [9].

A previous finite-element study incorporated region- and layer-specific FCD distributions to simulate Donnan swelling [10]. Osmotic effects contributed to opening but did not fully explain the regional depletion response. This discrepancy suggested that GAGs may also influence the locally preferred architecture of the wall through their interactions with the surrounding matrix.

We therefore tested a reduced mechanism in which the control wall contains a structural preferred-stretch pattern retained after GAG depletion together with a removable circumferential component shaped by the measured through-wall FCD distribution. The central question was whether the same coefficient governing this removable component could account for the regional mean depletion responses of the ascending aorta, arch, and descending aorta after each region’s control structural pattern had been fitted independently. The one-layer wall provided the primary parsimonious test, and a two-layer wall assessed anatomical robustness. We hypothesized that one FCD-linked coefficient would capture the three regional mean responses without requiring a separate depletion parameter for each region.

## 2 Methods

### 2.1 Experimental basis and model overview

The framework combined analytical axisymmetric kinematics and constitutive relations with numerical solution of radial equilibrium, opening-angle selection, and parameter fitting. It used regional-average porcine data from the ascending aorta, arch, and descending thoracic aorta. Closed-ring geometry and paired control/GAG-depleted circumferential opening angles were taken from Ghadie et al. [9]. Intramural sulfated-GAG distributions were taken from Ghadie et al. [8], converted to FCD using the conventions of Ghadie et al. [10], and represented as eight-bin regional mean profiles rather than specimen-specific profiles. The measured angles are listed in Table 1. Figure 1 summarizes the regional geometry, opening-angle convention, FCD profiles, and multiplicative stretch decomposition. Complete geometries, layer fractions, FCD bins, material parameters, and numerical constants are provided in Supplementary Tables S1–S7.

**Table 1:** Experimental circumferential opening angles used as direct model inputs [9]. Values are mean *±* SD from *N* = 5 porcine aortas, with adjacent control and GAG-depleted rings excised from each region. The lower descending thoracic region in the source study is denoted here as descending.

| Region | Control $\alpha$ | Depleted $\alpha$ | Reduction |
| --- | --- | --- | --- |
| Ascending | $114 \pm 17^\circ$ | $85 \pm 18^\circ$ | $29^\circ$ |
| Arch | $61 \pm 27^\circ$ | $41 \pm 24^\circ$ | $20^\circ$ |
| Descending | $31 \pm 9^\circ$ | $18 \pm 7^\circ$ | $13^\circ$ |

**Figure 1:**
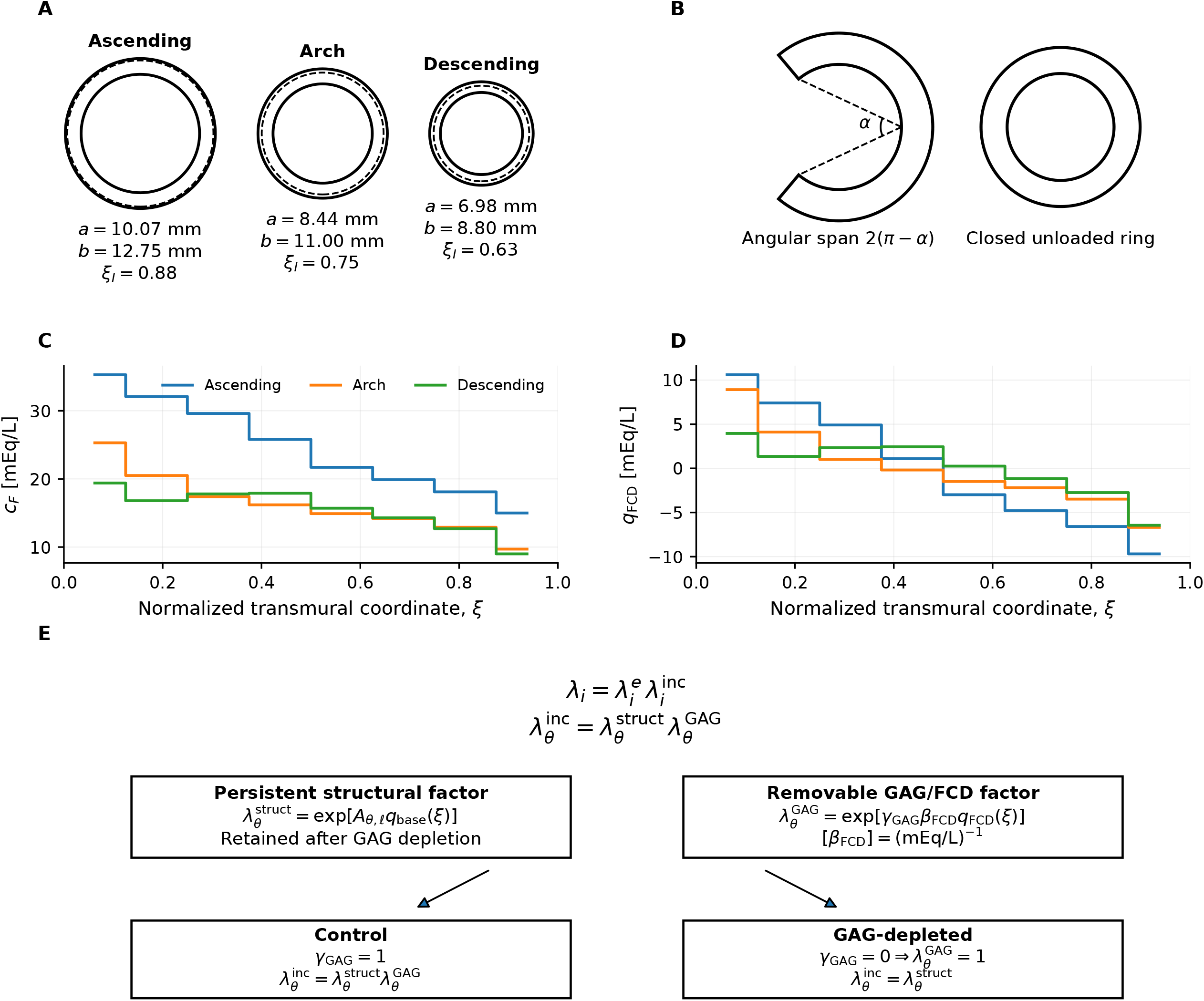
Regional geometry, experimental inputs, and incompatible-stretch decomposition. **(A)** Closed unloaded ring geometries; *a* and *b* are the inner and outer radii, and dashed circles denote the two-layer interfaces. **(B)** Opening-angle convention and closed unloaded ring; the measured regional control and depleted values are listed in Table 1. **(C)** Regional mean eight-bin FCD-magnitude profiles, where *c*_*F*_ is local FCD magnitude. **(D)** Regionally centered fields 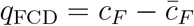. Panels C–D show regional mean profiles, not specimen-level profiles; variability in the original sulfated-GAG measurements was reported as SEM in the source study but was not propagated through the present regional-average model. **(E)** Multiplicative decomposition of total stretch into elastic and incompatible components, with the circumferential incompatible stretch further separated into a structural factor retained after depletion and a removable GAG/FCD-associated factor. Geometry and opening angles are from [9]; FCD profiles derive from [8] using the conversion and layer conventions of [10].

Published young-human longitudinal opening angles of 45°, 60°, and 50° for the ascending, arch, and descending regions were used only as soft targets to keep the fitted control axial field within a plausible range [11]. These values were neither species matched nor measured in the same specimens, and no corresponding GAG-depleted longitudinal measurements were available in the literature. The tolerance assigned to this axial check was *σ*_*ψ*_ = 15°, chosen from sensitivity analyses. Consequently, this term only discouraged extreme axial residual fields; *σ*_*ψ*_ was not treated as a measurement uncertainty.

The region-specific structural amplitudes were fitted only to the control state; the depleted opening angles were not used in that step. They were used afterward to ask whether one shared FCD-linked coefficient could account for the three mean responses to GAG depletion. Because only three regional mean depleted angles were available, the model was not intended to establish specimen-level predictivity or unique identifiability of all GAG-mediated mechanisms. Instead, it tested whether the observed regional mean depletion pattern could be represented without assigning an independent depletion parameter to each region.

### 2.2 Closure kinematics and incompatible stretch

An assumed stress-free sector with inner and outer radii *A* and *B*, coordinates (*R*, Θ, *Z*), and angular span 2(*π* − *α*) was closed into the measured ring (*r, θ, z*) using

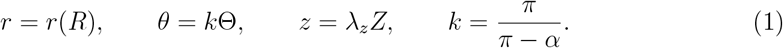

The total principal stretches were

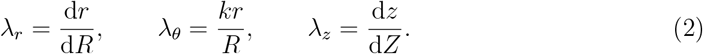

The model separates the stretch required to close the sector into two parts. The elastic part, 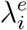, produces stress through the constitutive law. The preferred part, 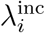, represents a local stress-free stretch that the tissue would adopt if that material point could deform independently. Because different wall locations are assigned different preferred stretches, these local stress-free states cannot generally be assembled into one closed ring without residual stress. In this sense the preferred-stretch field is incompatible. In the present model, the incompatible stretch was further decomposed into two multiplicative components: a structural preferred stretch retained after GAG depletion and a removable GAG-associated preferred stretch:

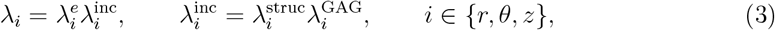

where 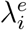 is the elastic stretch used in the constitutive law, 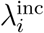 is the incompatible or preferred stretch, 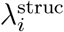 is the structural preferred stretch retained after GAG depletion, and 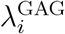 is the GAG-associated preferred stretch removed during depletion.

The centered linear structural shape was *q*_base_ = 2(*R* − *A*)*/*(*B* − *A*) − 1, so it varies from −1 at the inner wall to +1 at the outer wall. Equation 4 is a deliberately low-order model assumption for the retained transmural structural preferred-stretch gradient. For material layer £, the dimensionless logarithmic amplitudes *A*_*θ,f*_ and *A*_*z,f*_ controlled the circumferential and axial structural stretches:

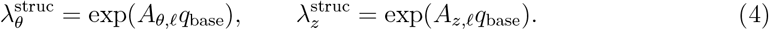

The removable FCD-linked field used 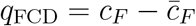, where *c*_*F*_ is the local FCD magnitude and 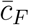 is its regional mean. Equation 5 is the proposed GAG/FCD-linked component of the present model. It was chosen as the simplest exponential/log-stretch relation in which the local removable preferred stretch is proportional to the centered FCD field. Centering makes the model sensitive to through-wall FCD variation rather than to a uniform regional offset. The coefficient *β*_FCD_ converts the local FCD departure into a circumferential logarithmic stretch:

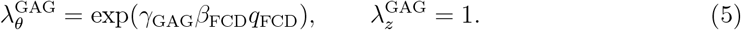

The scalar switch *γ*_GAG_ encoded whether the GAG-associated preferred-stretch component was present: *γ*_GAG_ = 1 in control tissue and *γ*_GAG_ = 0 after GAG depletion. Because Eqs. 4 and 5 define the structural and GAG/FCD-associated factors as exponentials, those stretch factors remain strictly positive for all fitted amplitudes; local volume preservation then gives a positive radial incompatible stretch through the reciprocal product of the circumferential and axial factors. The one-layer model used *A*_*θ*,base_ and *A*_*z*,base_ throughout the wall. The two-layer model used one circumferential amplitude, *A*_*θ*,shared_, in both layers and separate axial amplitudes *A*_*z,M*_ and *A*_*z,A*_ in the intima-media and adventitia. The circumferential amplitude was shared across layers to avoid giving the robustness model enough freedom to fit independent circumferential structural gradients in each layer while still allowing anatomical differentiation through the axial amplitudes. Full directional expressions are given in Supplementary Methods S2.

### 2.3 Constitutive response and equilibrium

The solid matrix followed the uncoupled Holmes–Mow law [12], as used in [10],

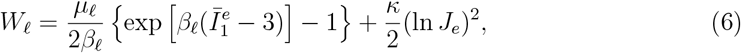

where 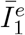 and *J*_*e*_ are computed from the elastic stretches. For each candidate opening angle, radial equilibrium and traction-free boundaries required

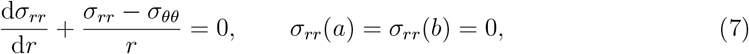

where *σ*_*rr*_ and *σ*_*θθ*_ are the radial and circumferential Cauchy stresses, respectively. The solution also enforced displacement and radial-stress continuity at the layer interface and zero resultant axial force, recovered the measured closed radii, and satisfied near-incompressibility and numerical-acceptance checks. Constitutive stresses, axial force, and complete boundary conditions are provided in Supplementary Methods S3.

### 2.4 Control calibration, depletion prediction, and joint coefficient selection

For each region and candidate *β*_FCD_, the structural amplitudes were identified from the control state only. These were *A*_*θ*,base_ and *A*_*z*,base_ in the one-layer model and *A*_*θ*,shared_, *A*_*z,M*_, and *A*_*z,A*_ in the two-layer model. The objective included the measured circumferential control angle, the soft longitudinal axial-field check, geometric closure, axial-force balance, volume-change diagnostics, and opening-angle selection checks. The depleted angle was not used in this step.

Depletion was simulated by changing *γ*_GAG_ from 1 to 0 while holding the fitted structural amplitudes fixed. Although 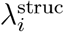 is held fixed during depletion, it is not irrelevant: it defines the residual-stretch architecture retained after removal of the GAG-associated component and therefore determines the elastic stretches and stresses obtained when the depleted ring is reclosed. Because numerical continuation and local angle searches can introduce small selection drift, each depletion calculation was paired with a no-depletion numerical control. In this control calculation, *γ*_GAG_ was kept equal to 1, so the GAG-associated preferred-stretch component remained present and no biological depletion was simulated. The selected angle from this run, *α*_no dep_, estimates the numerical drift associated with the angle-selection procedure. The corrected depleted prediction was

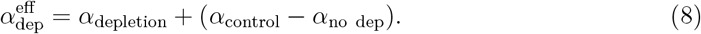

The magnitude |*α*_control_ − *α*_no dep_| was required to remain small relative to the predicted depletion effect; the corresponding values and acceptance checks are reported in Supplementary Methods S4 and Supplementary Figure S2A. One coefficient was selected jointly across the three regions by minimizing the equally weighted sum of the three regional residuals,

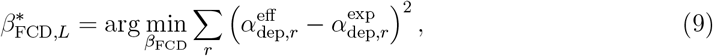

where *L* denotes the layer representation. The 5° residual scale used only to plot the nondimensional joint-error landscape in Figure 2 is reported in Supplementary Methods S5; it does not change the minimizing coefficient. Detailed calibration, continuation, and correction equations are given in Supplementary Methods S4–S5.

**Figure 2:**
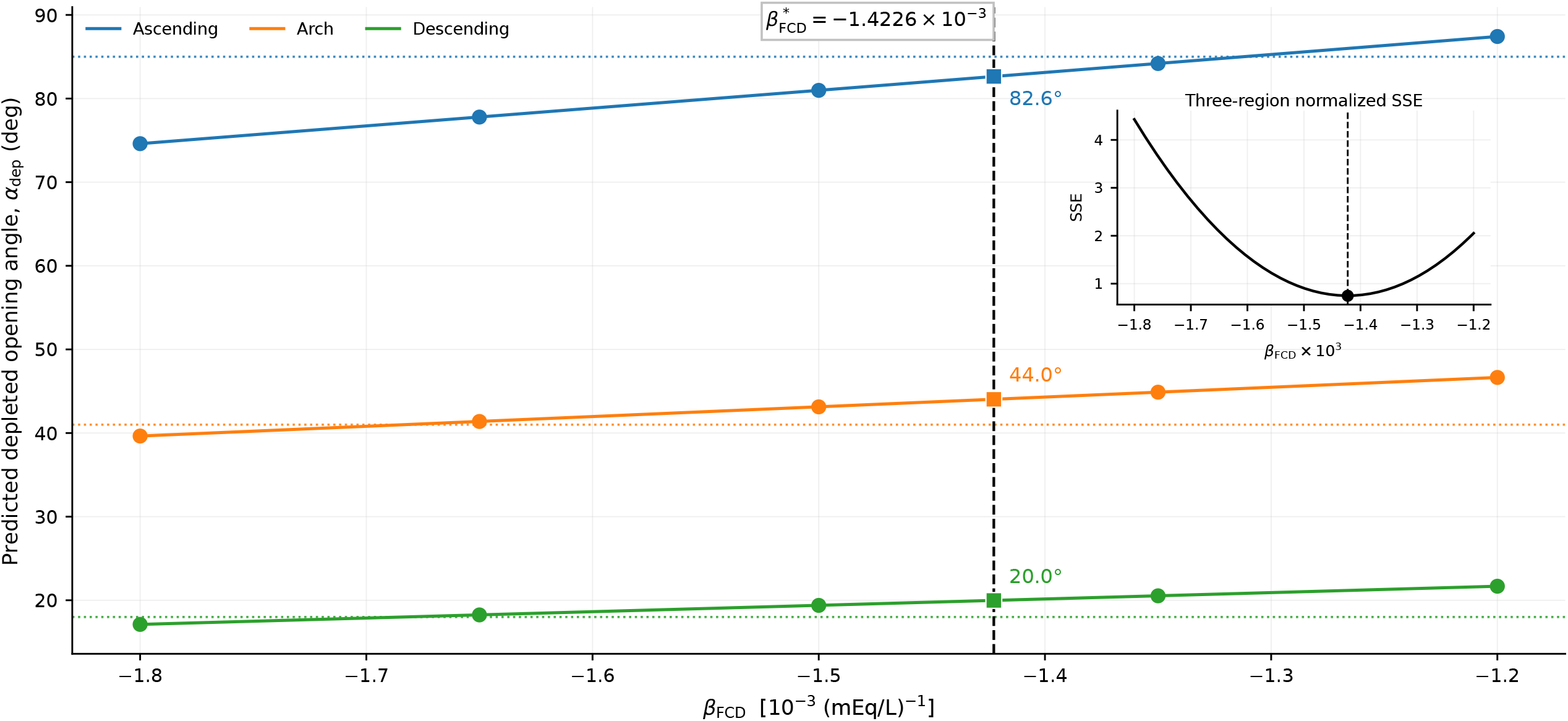
Selection of the shared FCD-linked coefficient in the primary one-layer model. Circles are evaluated points, curves are shape-preserving interpolants, dotted horizontal lines are measurements, and squares are predictions at the selected coefficient. The inset shows the joint normalized squared error used only for visualization of the coefficient-selection landscape. Measured-predicted agreement and signed errors are shown in Figure 3.

### 2.5 Testing osmotic pressure as the only depletion mechanism

We also asked whether the measured depletion response could be explained without introducing the FCD-linked preferred-stretch component. In this calculation, GAGs affected the wall only through the osmotic pressure implied by the local FCD magnitude. The FCD profile defined a local Donnan pressure *p*_*D*_(*R*); a dimensionless multiplier *χ*_*D*_ then scaled this contribution, 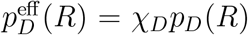. The resulting pressure entered the stress balance as an isotropic compressive contribution, 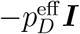. The same control-fitting and depletion-prediction workflow was then repeated, but without *λ*^GAG^. Depletion was represented by removing the FCD-derived pressure contribution rather than by removing an FCD-linked preferred stretch.

After this setup, we refer to the calculation as the Donnan-pressure-only comparator. A positive outcome would have been a predicted depleted opening angle that moved from the control value toward the measured depleted value after the osmotic contribution was removed. The multiplier *χ*_*D*_ was evaluated at 0, 0.5, and 1 for the ascending-aorta calibration; regional predictions were compared at the selected setting 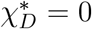. This was a three-point sensitivity analysis, not a continuous parameter sweep or a validation of osmotic swelling. The full pressure equation and constants are given in Supplementary Methods S6 and Table S6.

All computations used a custom implementation in MATLAB Version R2025b (The MathWorks, Inc., Natick, MA, USA). Primary outcomes were regional signed errors and the three-region RMSE; secondary outcomes were the selected coefficient, fitted structural amplitudes, no-depletion-control drift, numerical acceptance score, boundary flags, and longitudinal consistency. Here, 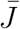 denotes the wall-averaged local volume ratio, so 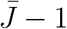 served as a scalar near-incompressibility diagnostic. Complete solver settings and acceptance criteria are given in Supplementary Table S7 and Supplementary Methods S7–S8.

## 3 Results

### 3.1 Osmotic pressure alone did not predict the depletion response

In the osmotic-pressure-only calculation, success would have meant that removing the FCD-derived pressure contribution produced a lower opening angle, moving the prediction from the control value toward the measured depleted value. This did not occur. In the ascending-aorta sensitivity analysis, the predicted depleted angle increased from 114° at *χ*_*D*_ = 0 to 119° and 124° at *χ*_*D*_ = 0.5 and 1, respectively, whereas the measured depleted target was 85°; the two nonzero settings also reached the prescribed structural-amplitude bound (Supplementary Figure S1A). Thus, increasing the Donnan-pressure contribution moved the prediction away from the experimental depletion response. At the selected setting, 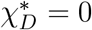, regional predictions remained at the control values of 114°, 61°, and 31°, compared with depleted measurements of 85°, 41°, and 18° (Supplementary Figure S1B). The Donnan-pressure-only comparator therefore failed as a sufficient explanation of the measured opening-angle reductions. This finding should not be interpreted as evidence that Donnan swelling is absent; rather, the imposed osmotic-pressure pathway alone did not provide the correct direction and magnitude of the depletion-induced opening-angle change.

### 3.2 One coefficient captured the three regional mean depletion responses

The one-layer analysis selected

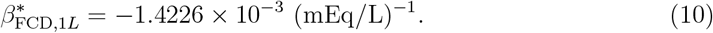

The regional calibration curves and joint-error landscape are shown in Figure 2. Predicted depleted angles were 82.639°, 44.032°, and 19.979° for the ascending, arch, and descending regions, with signed errors of −2.361°, +3.032°, and +1.979°. The three-region RMSE was 2.50°, the mean absolute error was 2.46°, and the maximum absolute error was 3.03°.

The two-layer analysis selected

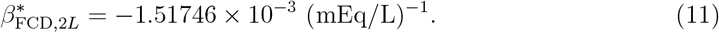

Predicted angles were 82.972°, 44.273°, and 18.534°, with signed errors of −2.028°, +3.273°, and +0.534°. The RMSE was 2.24° and the mean absolute error was 1.94°. Table 2 and Figure 3 compare the models. Layer differentiation improved the descending prediction but did not reduce the largest regional error.

**Table 2:** Measured and predicted depleted opening angles. Parentheses give signed error, predicted minus measured.

| Region | Measured<br>[°] | One-layer<br>[°] | Two-layer<br>[°] |
| --- | --- | --- | --- |
| Ascending | 85 | 82.639 (−2.361) | 82.972 (−2.028) |
| Arch | 41 | 44.032 (+3.032) | 44.273 (+3.273) |
| Descending | 18 | 19.979 (+1.979) | 18.534 (+0.534) |

**Figure 3:**
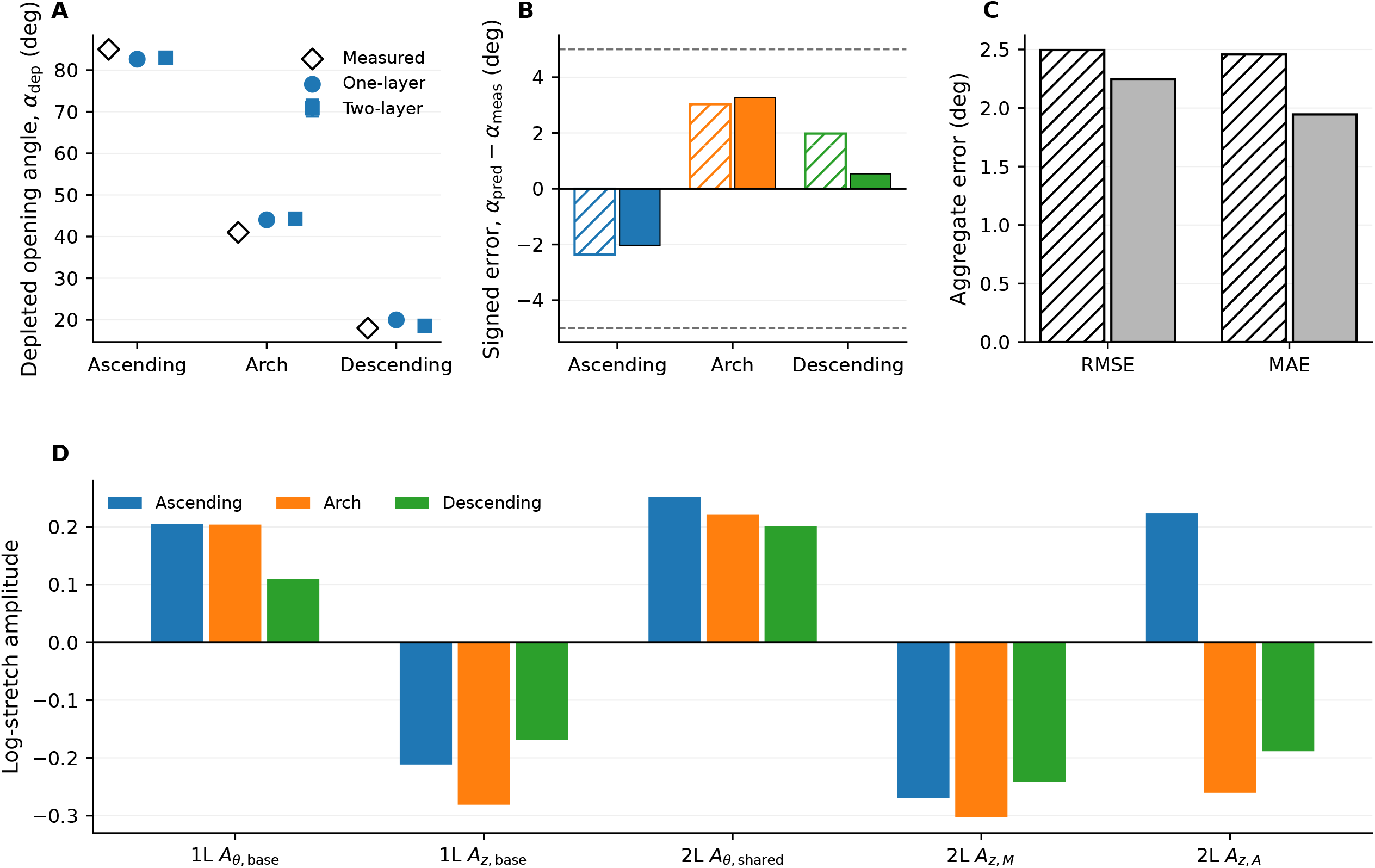
Comparison of the primary one-layer model and the two-layer robustness model. **(A)** Regional measured and predicted depleted angles. **(B)** Signed regional prediction errors; hatched bars denote the one-layer model and solid bars denote the two-layer model. **(C)** Aggregate RMSE and mean absolute error, using the same model styles as in panel B. **(D)** Fitted structural amplitudes for the one- and two-layer models, expressed as dimensionless logarithmic-stretch coefficients multiplying the structural shape function *q*_base_ defined in Section 2.2.

### 3.3 Layer differentiation did not create the mechanism

The two fitted coefficients differed by only 6.7% in magnitude, and both models reproduced every regional depleted angle within approximately 3.3° (Figure 3A–C). The one-layer model used two structural amplitudes per region; the two-layer model used one shared circumferential amplitude and two layer-specific axial amplitudes. The circumferential amplitude was positive in every region and model, and the one-layer or medial axial amplitude was negative in every region. The adventitial axial amplitude changed sign, indicating weaker transferability (Figure 3D; Supplementary Table S8).

The longitudinal calculations were retained only as checks on the fitted axial fields. Predicted control longitudinal angles were within 0.27° of the published longitudinal reference values because the objective included them as a soft axial-field check. Depletion changed the predicted longitudinal angle by −1.08° to +0.07° in the one-layer model and −0.89° to −0.11° in the two-layer model (Supplementary Figure S3), but these small changes should not be interpreted as validated axial predictions. All final cases passed the opening-angle-window and transfer-edge checks; the maximum 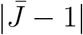 was 1.73 *×* 10^−4^ and the maximum radius mismatch was 0.27 *µ*m (Supplementary Figure S2).

## 4 Discussion

### 4.1 Principal finding and biological interpretation

A single FCD-linked coefficient captured the regional mean depleted circumferential opening angles of all three regions within 3.1° in the primary one-layer model (RMSE 2.50°). Introducing layer-specific axial amplitudes reduced the RMSE only to 2.24°. The central result is therefore not that layer differentiation is necessary, but that the three regional mean reductions did not require a separate GAG-depletion parameter for each region. After the control state was matched region by region, the depletion response was captured by one shared FCD-linked coefficient. This does not independently validate a predictive model, because the same three regional mean depleted angles were used to select and assess the shared coefficient.

This mechanism complements rather than replaces Donnan swelling. In the previous finite-element study, FCD-derived swelling was evaluated in terms of the experimental decrease in opening angle caused by GAG removal, denoted “Delta” in that study and defined as the difference between control and GAG-depleted opening angles [10]. In that sense, swelling was interpreted as contributing to the part of the control opening angle lost after depletion, rather than as recovering the full control opening angle by itself. The present osmotic-pressure-only calculation was included as a mechanistic counterfactual: if osmotic pressure were sufficient to explain the depletion response, then removing the FCD-derived pressure contribution should have reduced the predicted opening angle toward the measured depleted values. Instead, nonzero Donnan-pressure settings moved the ascending prediction away from the depleted target, and the best regional comparison occurred when the Donnan-pressure multiplier was zero. This result should not be read as evidence that Donnan swelling is absent or biologically irrelevant. It means that, in this closure framework, the imposed Donnan-pressure term alone did not provide the correct direction and magnitude of the depletion-induced opening-angle change. The FCD-shaped preferred stretch may summarize GAG-mediated effects that opening-angle data cannot separate, including osmotic prestress, interactions with elastin and collagen, and altered fiber recruitment after depletion [6, 7]. A coupled Donnan-plus-incompatible-stretch model is plausible, but both effects follow the same FCD distribution and are removed together experimentally, so their amplitudes are not identifiable from the present observations alone.

### 4.2 Parsimony, layer structure, and material dependence

The one-layer model is preferred because it achieves nearly the same accuracy with fewer independently fitted structural amplitudes. The two-layer result remains valuable: it shows that anatomical differentiation preserves the selected coefficient and regional predictions rather than creating the effect. The shared circumferential and medial axial patterns appear more robust than the adventitial axial amplitude.

Supplementary Figure S2 shows that the representative total closure stretches remained close to unity and that the associated Cauchy stresses were on the order of several kilopascals. The calculations therefore remained relatively near the reference state, where constitutive models with comparable near-reference tangent stiffness would be expected to produce similar incremental depletion responses. This observation supports, but does not establish, qualitative robustness to the material law: the constitutive law acts on elastic rather than total stretches, and differences in anisotropy, collagen recruitment, and layer behavior may still alter the fitted residual fields and detailed regional predictions.

The Holmes–Mow law was retained for consistency with [10] and because its uncoupled form is available in FEBio [13, 14]. It does not encode explicit collagen directions or layer-specific anisotropy. Testing an anisotropic arterial model is therefore required before claiming quantitative material-law independence.

### 4.3 Interpretation, reuse, and pathological extension

Because *β*_FCD_ multiplies the centered field *q*_FCD_, the mechanically relevant quantity is the dimensionless product *β*_FCD_*q*_FCD_. Centering isolates through-wall variation and excludes a uniform regional FCD contribution from this incompatible-stretch mechanism. The similar one- and two-layer coefficients support a stable mechanism-level estimate, but not a universal biological constant.

The fitted coefficient and structural amplitudes can be reused directly only when the same FCD centering, structural shape function, opening-angle convention, constitutive law, and regional-average geometry are retained. Otherwise they are better treated as informed initial estimates. Using one coefficient here does not restrict extension to aneurysmal or remodeled tissue: pathological geometry, material properties, FCD distributions, and retained structural fields can be introduced within the same formulation. Whether *β*_FCD_ itself transfers across healthy and pathological states must be tested; disease-specific or hierarchical coefficients can be introduced when appropriate data become available.

### 4.4 Limitations and future validation

The model is reduced and axisymmetric and is intended to compare mechanisms rather than replace three-dimensional finite-element analysis. Structural amplitudes are fitted parameters and are not uniquely identifiable from opening angles alone. The same three regional depleted angles were used to select and assess the shared coefficient, so the results demonstrate cross-regional representation rather than external validation. The present analysis used regional-average inputs and outputs; it therefore does not test whether the same coefficient predicts specimen-to-specimen variation. Such a test would require paired specimen-specific opening angles, specimen-specific geometry, local FCD profiles, and preferably local strain or stress-related measurements.

Paired depleted longitudinal angles were unavailable, and the predominantly circumferential response remains unvalidated in that direction. The axial amplitudes, especially *A*_*z,A*_, should be interpreted cautiously: their role was to prevent implausible axial residual fields during circumferential calibration, not to predict a validated longitudinal depletion response. Donnan and incompatible-stretch contributions cannot be separated uniquely with the present intervention. The results also depend on the centered eight-bin FCD profiles and Holmes– Mow material representation. Independent specimens, local strain measurements, paired circumferential and longitudinal depletion data, and alternative constitutive laws are the next requirements for predictive validation.

## 5 Conclusions

This study introduced a reduced closure framework that separated control residual architecture into a structural field retained after GAG removal and a removable circumferential component shaped by the FCD profile. One coefficient captured the regional mean depleted opening angles of three porcine thoracic aortic regions with a one-layer RMSE of 2.50°; the two-layer model reduced the RMSE only modestly to 2.24° and changed the coefficient by 6.7%.

The one-layer model is therefore the preferred parsimonious representation, while the two-layer model provides anatomical robustness. The findings support a role for GAGs in aortic residual mechanics that extends beyond Donnan pressure alone. Independent specimen-level FCD profiles, geometry, opening angles, and longitudinal depletion data are needed for predictive validation.

## Acknowledgments

The authors acknowledge the computational and editorial assistance of OpenAI’s ChatGPT, as detailed in the following declaration. All scientific decisions, model assumptions, computations, source verification, interpretations, and final wording were independently reviewed and approved by the authors.

## Declaration of generative AI and AI-assisted technologies in manuscript preparation

During preparation of this work, the authors used OpenAI’s ChatGPT to assist with manuscript organization and revision, mathematical exposition, computer-code development and debugging, literature searching, and exploration of alternative modeling and numerical-analysis strategies. The authors critically reviewed and, where necessary, independently verified all generated material and take full responsibility for the published content.

## Notes

### Competing Interest Statement

The authors have declared no competing interest.

### Summary of Updates

The revision substantially improved clarity, scope, and transparency without changing the underlying analyses or principal conclusions. The manuscript now presents the work more cautiously. Rather than claiming that the model explains the depletion response, it states that one shared FCD-linked coefficient can capture the mean regional response after each control state is fitted independently. It also makes clear that this is not specimen-level validation or unique identification of the underlying mechanism. The Methods were rewritten to explain the stretch decomposition, the meaning of preferred or incompatible stretch, the structural and GAG-associated components, and the switch used to represent GAG presence or depletion. The role of the structural amplitudes and the rationale for the two-layer formulation were also clarified. Longitudinal opening-angle information is now described as a soft constraint used to discourage implausible axial fields, rather than with specialized terminology. Longitudinal outputs are treated as consistency checks rather than validated predictions. The provenance of the FCD profiles was clarified. They are regional-average profiles derived from previously measured transmural sulfated GAG distributions, not specimen-specific data. Table 1 now also includes experimental variability and sample size for the control and GAG-depleted opening-angle measurements. The no-depletion correction and the coefficient-selection procedure were rewritten in more accessible terms. The Donnan analysis was also substantially improved by explaining what physical question is being tested, what a successful pressure-only outcome would have looked like, and why the actual result is insufficient. The conclusion is that the imposed Donnan-pressure pathway alone cannot reproduce the depletion response, not that Donnan swelling is absent. The Discussion now distinguishes more clearly between fitting, testing, prediction, and mechanistic interpretation, and states the study limitations more explicitly. Figures were streamlined to reduce redundancy, the Supplementary Information was refocused on reproducibility rather than repetition, terminology was simplified throughout, and the references were corrected and reordered.

