## Supplemental information for "A reduced glycosaminoglycan-linked residual-strain model captures regional opening angle changes after depletion in the porcine thoracic aorta"

**3 Supplementary Figures** **8**

### 1 Supplementary Methods

#### S1. Regional inputs and FCD conversion

The regional closed-ring geometry, layer fractions, opening-angle targets, FCD profiles, material constants, and numerical settings are listed in Supplementary Tables S1–S7. Lengths were converted to SI units internally. This section is intended as a reproducibility record rather than a repetition of the conceptual model described in the main text. The FCD magnitudes were derived from regional mean serial 300 $\mu$m cryosections quantified by dimethylmethylene-blue assay [1]. Those source measurements were reported with SEM, but the present reduced model used only the regional mean FCD profiles and did not propagate FCD uncertainty. The conversion in [2] used two negative charges per chondroitin-sulfate molecule, molecular weight 513 g/mol, and wall water fraction 0.70 to obtain signed FCD from sulfated-GAG mass. The present model used the magnitude $c_F$ and centered it within each region,

$$q_{\mathrm{FCD},j} = c_{F,j} - \bar{c}_F, \qquad \bar{c}_F = \frac{1}{8}\sum_{j=1}^{8} c_{F,j}. \tag{S1}$$

The field was centered but not standardized, so $q_{\mathrm{FCD}}$ retains units of mEq/L.

Table S1: Closed-ring geometry. $D_o = 2b$ is the closed unloaded outer diameter.

| Region | $D_o$ [mm] | $h$ [mm] | $a$ [mm] | $b$ [mm] | $H_{\mathrm{fit}}$ [mm] |
|---|---|---|---|---|---|
| Ascending | 25.50 | 2.68 | 10.07 | 12.75 | 2.00 |
| Arch | 22.00 | 2.56 | 8.44 | 11.00 | 1.96 |
| Descending | 17.60 | 1.82 | 6.98 | 8.80 | 1.40 |

Table S2: Layer fractions and opening-angle targets.

| Region | $\xi_I$ | Layer assignment | $\alpha_{\mathrm{ctl}}$ | $\alpha_{\mathrm{dep}}$ | $\Delta\alpha$ |
|---|---|---|---|---|---|
| Ascending | 0.88 | 7/8 intima-media | 114° | 85° | 29° |
| Arch | 0.75 | 6/8 intima-media | 61° | 41° | 20° |
| Descending | 0.63 | 5/8 intima-media | 31° | 18° | 13° |

Table S3: Eight-bin FCD magnitudes [mEq/L], ordered from inner to outer wall.

| Region | $c_{F,j}$, $j = 1, \ldots, 8$ |
|---|---|
| Ascending | 35.3, 32.1, 29.6, 25.8, 21.7, 19.9, 18.1, 15.0 |
| Arch | 25.3, 20.5, 17.4, 16.2, 14.9, 14.2, 12.9, 9.7 |
| Descending | 19.4, 16.8, 17.8, 17.9, 15.7, 14.3, 12.7, 9.0 |

Table S4: One-layer uncoupled Holmes–Mow constants. $\mu_{\mathrm{mem}}$ is the membrane shear-stiffness parameter obtained from biaxial fitting; the volumetric shear modulus $\mu$ used in the closure calculation was obtained by dividing $\mu_{\mathrm{mem}}$ by the corresponding fitted specimen thickness $H_{\mathrm{fit}}$.

| Region | $\mu_{\mathrm{mem}}$ [N/m] | $\beta$ | $\mu$ [kPa] |
|---|---|---|---|
| Ascending | 61.31 | 2.09 | 30.66 |
| Arch | 65.99 | 2.56 | 33.67 |
| Descending | 58.90 | 2.37 | 42.07 |

Table S5: Two-layer uncoupled Holmes–Mow constants; M denotes intima-media and A adventitia. $\mu_{M,\mathrm{mem}}$ and $\mu_{A,\mathrm{mem}}$ are membrane shear-stiffness parameters obtained from biaxial fitting; $\mu_M$ and $\mu_A$ are the corresponding volumetric shear moduli used in the closure calculation after division by $H_{\mathrm{fit}}$.

| Region | $\mu_A/\mu_M$ | $\mu_{M,\mathrm{mem}}$ [N/m] | $\mu_{A,\mathrm{mem}}$ [N/m] | $\beta_M$ | $\beta_A$ | $\mu_M$ [kPa] | $\mu_A$ [kPa] |
|---|---|---|---|---|---|---|---|
| Ascending | 0.10 | 55.65 | 5.565 | 2.00 | 2.77 | 27.83 | 2.78 |
| Arch | 0.50 | 43.30 | 21.65 | 2.90 | 1.82 | 22.09 | 11.05 |
| Descending | 1.00 | 28.60 | 28.60 | 2.98 | 1.78 | 20.43 | 20.43 |

Table S6: Donnan and near-incompressibility constants.

| Quantity | Value | Meaning |
|---|---|---|
| $\phi_0$ | 0.70 | Reference fluid volume fraction |
| $c^*$ | 300 | External bath concentration scale |
| $R$ | 8.314462618 | Universal gas constant [J mol$^{-1}$ K$^{-1}$] |
| $T$ | 310 | Absolute temperature [K] |
| $\Phi$ | 1.0 | Osmotic coefficient |
| $\chi_D$ | 0, 0.5, 1.0 | Values in sensitivity analysis |
| $\kappa$ | $2.0 \times 10^6$ | Volumetric penalty [Pa] |

Table S7: Numerical constants.

| Quantity | Value | Meaning |
|---|---|---|
| $\lambda_z$ | 1.0 | Fixed axial stretch in reported joint regional runs |
| $\lambda_z$ bounds | [0.85, 1.15] | Bounds when allowed to vary |
| $\lambda_r$ bounds | [0.03, 30.0] | Admissible radial stretch interval |
| $J_{\min}$ | $10^{-6}$ | Minimum admissible local volume ratio |
| $\sigma_\alpha$ | $5^\circ$ | Circumferential residual scaling |

| Quantity | Value | Meaning |
|---|---|---|
| $\sigma_\psi$ | 15° | Tolerance for soft axial-field check |
| Amplitude bounds | [-0.50, 0.50] | Logarithmic-stretch amplitudes |
| Radial mesh | 81 | BVP mesh points |
| Evaluation points | 251 | Output and diagnostic points |
| BVP relative tolerance | $10^{-5}$ | Solver tolerance |
| BVP absolute tolerance | $10^{-8}$ | Solver tolerance |
| Radius weight | $10^{6}$ | Radius-mismatch objective weight |
| Axial-force weight | 1.0 | Axial-force residual weight |
| Volume-diagnostic weight | $10^{-2}$ | Mean volume-change diagnostic weight |
| Amplitude penalty weight | $10^{-4}$ | Small-amplitude penalty |

#### S2. Closure kinematics and incompatible-stretch decomposition

The opening sector used $\Theta \in [-(\pi-\alpha), \pi-\alpha]$ and $k = \pi/(\pi-\alpha)$. The deformation and total stretches were

$$r = r(R), \qquad \theta = k\Theta, \qquad z = \lambda_z Z, \tag{S2}$$

$$\lambda_r = \frac{\mathrm{d}r}{\mathrm{d}R}, \qquad \lambda_\theta = \frac{kr}{R}, \qquad \lambda_z = \frac{\mathrm{d}z}{\mathrm{d}Z}. \tag{S3}$$

The area relation was $(\pi-\alpha)(B^2 - A^2) = \pi(b^2 - a^2)$. For implementation, the elastic stretches used in the constitutive law were computed as

$$\lambda_i^e = \frac{\lambda_i}{\lambda_i^{\mathrm{struc}}\lambda_i^{\mathrm{GAG}}}. \tag{S4}$$

With $q_{\mathrm{base}} = 2(R-A)/(B-A) - 1$,

$$\lambda_\theta^{\mathrm{struc}} = \exp(A_{\theta,\ell} q_{\mathrm{base}}), \qquad \lambda_z^{\mathrm{struc}} = \exp(A_{z,\ell} q_{\mathrm{base}}), \tag{S5}$$

$$\lambda_\theta^{\mathrm{GAG}} = \exp(\gamma_{\mathrm{GAG}}\beta_{\mathrm{FCD}} q_{\mathrm{FCD}}), \qquad \lambda_z^{\mathrm{GAG}} = 1. \tag{S6}$$

The directional elastic stretches used in the computation were therefore:

$$\lambda_r^e = \frac{\mathrm{d}r}{\mathrm{d}R} \exp[(A_{\theta,\ell} + A_{z,\ell}) q_{\mathrm{base}} + \gamma_{\mathrm{GAG}}\beta_{\mathrm{FCD}} q_{\mathrm{FCD}}], \tag{S7}$$

$$\lambda_\theta^e = \frac{kr}{R} \exp[-A_{\theta,\ell} q_{\mathrm{base}} - \gamma_{\mathrm{GAG}}\beta_{\mathrm{FCD}} q_{\mathrm{FCD}}], \tag{S8}$$

$$\lambda_z^e = \lambda_z \exp[-A_{z,\ell} q_{\mathrm{base}}]. \tag{S9}$$

#### S3. Constitutive stress, equilibrium, and boundary conditions

The elastic tensors were $\boldsymbol{C}_e = \boldsymbol{F}_e^T \boldsymbol{F}_e$, $\boldsymbol{B}_e = \boldsymbol{F}_e \boldsymbol{F}_e^T$, $J_e = \det \boldsymbol{F}_e$, and $\bar{\boldsymbol{B}}_e = J_e^{-2/3} \boldsymbol{B}_e$. The uncoupled Holmes–Mow energy was

$$W_\ell = \frac{\mu_\ell}{2\beta_\ell}\{\exp[\beta_\ell(\bar{I}_1^e - 3)] - 1\} + \frac{\kappa}{2}(\ln J_e)^2, \tag{S10}$$

with Cauchy stress

$$\boldsymbol{\sigma}_\ell = \frac{\mu_\ell}{J_e}\exp[\beta_\ell(\bar{I}_1^e - 3)]\,\mathrm{dev}(\bar{\boldsymbol{B}}_e) + \frac{\kappa \ln J_e}{J_e}\boldsymbol{I}. \tag{S11}$$

Radial equilibrium and boundaries were

$$\frac{\mathrm{d}\sigma_{rr}}{\mathrm{d}r} + \frac{\sigma_{rr} - \sigma_{\theta\theta}}{r} = 0, \qquad \sigma_{rr}(a) = \sigma_{rr}(b) = 0, \tag{S12}$$

where $\sigma_{rr}$ and $\sigma_{\theta\theta}$ are the radial and circumferential Cauchy stresses, respectively. Two-layer cases also enforced continuity at the layer interface and the zero axial resultant condition

$$F_z = 2\pi \int_a^b \sigma_{zz} r \,\mathrm{d}r = 0. \tag{S13}$$

#### S4. Control calibration and no-depletion numerical correction

For each region, layer model, and candidate $\beta_{\mathrm{FCD}}$, the structural amplitudes minimized a control objective containing the circumferential mismatch, soft longitudinal axial-field check, radius mismatch, axial force, volume-change diagnostics, and opening-angle selection checks. The depleted angle was excluded from this objective. The computation followed this sequence:

1. Fit structural amplitudes in the control state with $\gamma_{\mathrm{GAG}} = 1$, representing the presence of the GAG-associated preferred-stretch component.
2. Resolve closure and select the control opening angle using the local angle-search procedure.
3. Repeat the same numerical continuation with $\gamma_{\mathrm{GAG}} = 1$ to keep the GAG-associated component present and obtain the no-depletion numerical control $\alpha_{\mathrm{no\ dep}}$.
4. Set $\gamma_{\mathrm{GAG}} = 0$ to remove the GAG-associated component, keep the fitted structural amplitudes fixed, resolve closure, and obtain $\alpha_{\mathrm{depletion}}$.
5. Correct the depletion prediction for angle-selection drift using

$$\alpha_{\mathrm{dep}}^{\mathrm{eff}} = \alpha_{\mathrm{depletion}} + (\alpha_{\mathrm{control}} - \alpha_{\mathrm{no\ dep}}). \tag{S14}$$

The numerical drift diagnostic was $\delta_{\mathrm{no\ dep}} = \alpha_{\mathrm{no\ dep}} - \alpha_{\mathrm{control}}$.

#### S5. Joint regional selection of the FCD-linked coefficient

At each candidate coefficient, the control structural amplitudes were re-identified independently in all three regions. The shared coefficient was selected by minimizing the equally weighted regional residual sum of squares,

$$\beta^*_{\mathrm{FCD},L} = \arg\min_{\beta_{\mathrm{FCD}}} \sum_r \left(\alpha^{\mathrm{eff}}_{\mathrm{dep},r} - \alpha^{\mathrm{exp}}_{\mathrm{dep},r}\right)^2. \tag{S15}$$

For plotting the coefficient-selection landscape, the same residuals were also divided by 5° to give a nondimensional normalized SSE. Because this was a common positive scale applied to all regions, it did not alter the minimizing coefficient; it served only as a reporting scale and was not an experimental standard deviation.

#### S6. Osmotic-pressure-only comparison

This comparison tested the specific hypothesis that removal of an FCD-derived osmotic pressure contribution is sufficient to reproduce the measured depletion-induced opening-angle reductions. In this calculation, no FCD-dependent preferred-stretch factor was included; equivalently, $\lambda_\theta^{\mathrm{GAG}} = 1$ throughout the wall. GAGs entered only through an isotropic pressure contribution derived from the local FCD magnitude.

For a monovalent salt bath, the local Donnan pressure was computed as

$$p_D(R) = RT\left[\sqrt{c_F(R)^2 + 4c_s^2} - 2c_s\right], \qquad p_D^{\mathrm{eff}}(R) = \chi_D p_D(R), \tag{S16}$$

where $R$ is the gas constant, $T$ is absolute temperature, $c_F(R)$ is the local FCD magnitude expressed in concentration units consistent with the bath salt concentration $c_s$, and $\chi_D$ is a dimensionless scaling multiplier. The term in brackets is the difference between the mobile-ion concentration implied inside the charged tissue and the corresponding bath value. Multiplication by $RT$ converts this concentration difference into an osmotic pressure. The effective pressure contributed $-p_D^{\mathrm{eff}}\boldsymbol{I}$ to the Cauchy stress; the negative sign denotes an isotropic compressive pressure contribution in the solid stress convention used here.

The control state was calibrated with this pressure contribution included, and the depletion prediction was obtained by removing the FCD-derived pressure contribution while keeping the fitted structural amplitudes fixed. A successful osmotic-pressure-only explanation would therefore have produced a depleted opening angle lower than the control angle and closer to the measured depleted angle. Instead, the ascending-aorta sensitivity analysis at $\chi_D = 0$, 0.5, and 1 showed that the nonzero settings moved the prediction away from the measured depleted value. Regional comparisons therefore used the selected setting $\chi_D^* = 0$. This was a three-point sensitivity comparison, not a continuous calibration of Donnan pressure and not a test of whether Donnan swelling exists biologically.

##### S7. Longitudinal checks

The longitudinal release calculation used the same residual field and material parameters with the longitudinal release geometry. No depleted longitudinal target was used. Because $\lambda_z^{\mathrm{GAG}} = 1$, depletion had only an indirect predicted effect on longitudinal opening.

##### S8. Numerical acceptance criteria

Recorded diagnostics included no-depletion-control drift, radius mismatches, $\bar{J} - 1$, axial-force residual, the numerical acceptance score, and whether the selected angle occurred at a search-window boundary. Here $\bar{J}$ denotes the wall-averaged local volume ratio. Solutions were rejected or flagged when geometric closure failed, diagnostic tolerances were exceeded, or the selected angle lay at the search-window edge.

#### 2 Supplementary Results

##### S9. Fitted structural amplitudes

Table S8: Fitted structural amplitudes at the selected coefficients. Entries are dimensionless logarithmic-stretch amplitudes multiplying $q_{\mathrm{base}}$.

| | One-layer | | Two-layer | | |
|---|---|---|---|---|---|
| Region | $A_{\theta,\mathrm{base}}$ | $A_{z,\mathrm{base}}$ | $A_{\theta,\mathrm{shared}}$ | $A_{z,M}$ | $A_{z,A}$ |
| Ascending | 0.205042 | -0.211557 | 0.252654 | -0.269868 | 0.223376 |
| Arch | 0.204036 | -0.281150 | 0.221008 | -0.302687 | -0.260565 |
| Descending | 0.110253 | -0.168933 | 0.201375 | -0.241133 | -0.188397 |

#### 3 Supplementary Figures

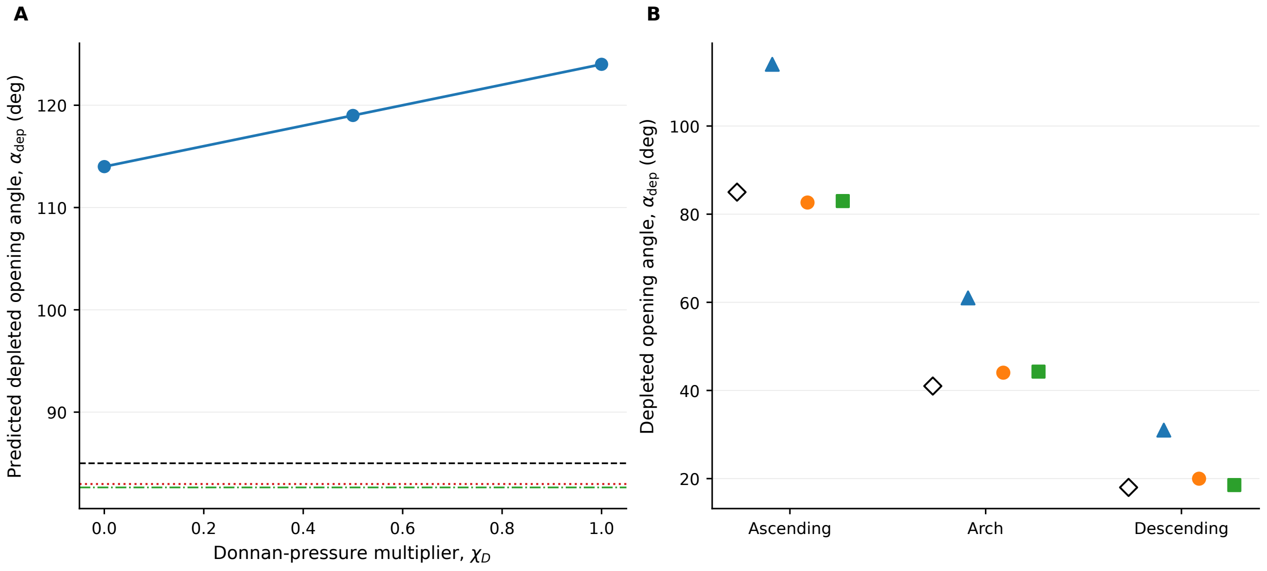

Figure S1: Osmotic-pressure-only comparison. **(A)** Ascending-aorta three-point sensitivity analysis over $\chi_D = 0$–1. The calculation asks whether removing the FCD-derived pressure contribution alone can move the predicted depleted angle from the control value toward the measured depleted value. The blue curve shows the effective depleted angle after control-state recalibration, and the horizontal lines show the measured target and the one- and two-layer preferred-stretch predictions. The $\chi_D = 0.5$ and 1 calibrations reached the prescribed structural-amplitude bound and are interpreted as boundary-limited sensitivity results. **(B)** Regional mechanism comparison at the selected setting $\chi_D^* = 0$, showing measured angles and the osmotic-pressure-only, one-layer, and two-layer predictions. The sensitivity analysis was performed for the ascending calibration; the regional comparison was evaluated at the selected setting.

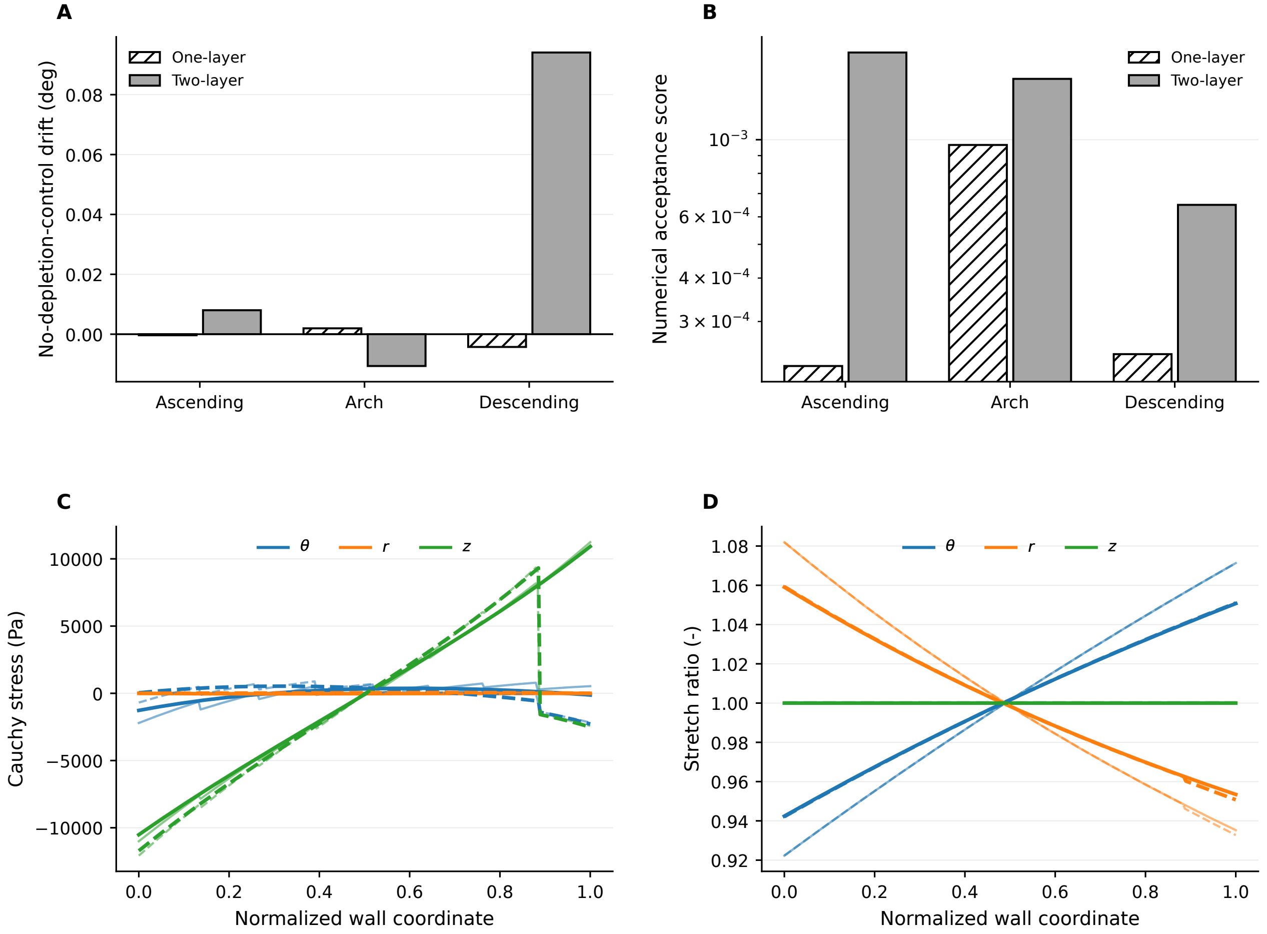

Figure S2: Numerical diagnostics and representative through-wall fields. **(A)** No-depletion-control drift for the three regions and both layer representations. **(B)** Numerical acceptance scores for the same cases. **(C)** Ascending-aorta Cauchy stresses before and after depletion; color distinguishes $\theta$, $r$, and $z$, solid and dashed lines denote one- and two-layer models, and lighter/thinner and darker/thicker curves denote control and depleted states. **(D)** Ascending-aorta stretch ratios before and after depletion, using the same conventions.

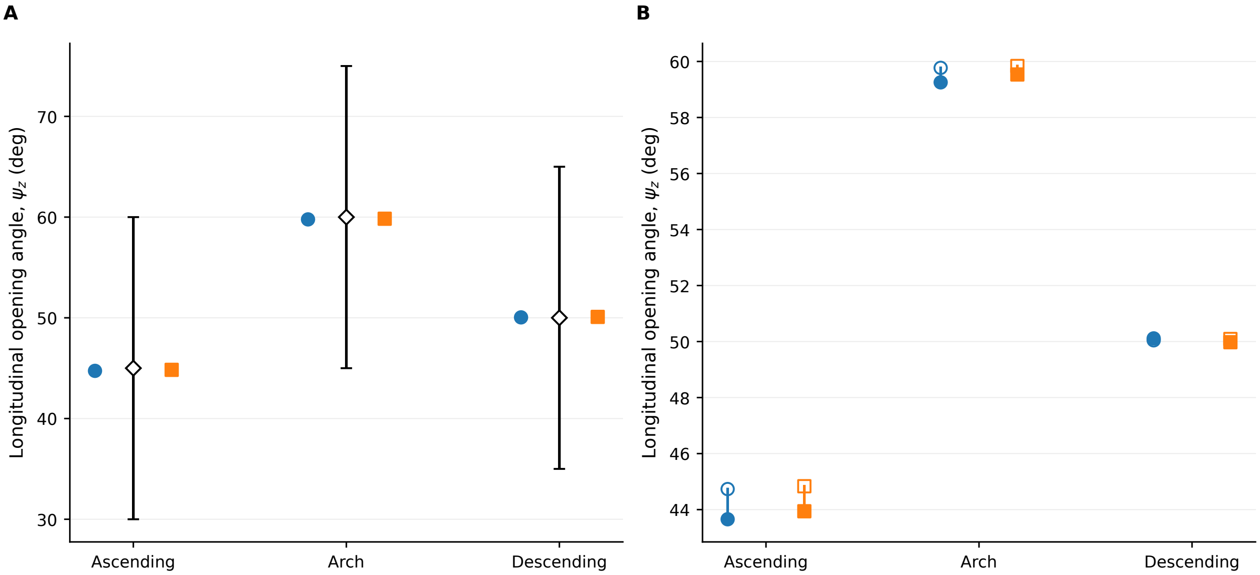

Figure S3: Longitudinal checks on fitted axial fields. **(A)** Control-state comparison of published young-human references (open diamonds), shown with the sensitivity-informed ±15° tolerance band $\sigma_\psi = 15°$, and the one- and two-layer control predictions. **(B)** Model-predicted control and depleted longitudinal opening angles for the same regions. Open diamonds denote published references; filled circles and squares denote one- and two-layer predictions, respectively. No depleted longitudinal target was used in calibration or coefficient selection.
